# Aberrant preparation of hand movement in schizophrenia spectrum disorder: An fMRI study

**DOI:** 10.1101/2025.03.23.644290

**Authors:** Harun A. Rashid, Tilo Kircher, Benjamin Straube

## Abstract

Schizophrenia spectrum disorder (SSD) is linked to impaired self-other distinction and action feedback monitoring, largely stemming from sensory-motor predictive mechanisms. However, the neural correlates of these predictive processes during movement preparation are unknown. Here, we investigated whether patients with SSD exhibit aberrant sensory-motor predictive processes reflected in neural activation patterns prior to hand movement onset. Functional MRI data from patients with SSD (n = 20) and healthy controls (n = 20) were acquired during actively performed or passively induced hand movements. The task required participants to detect temporal delays between their movements and video feedback, which either displayed their own (self) or someone else’s (other) hand moving in accordance with their own hand movements.

Patients compared to healthy controls showed reduced preparatory blood-oxygen-level-dependent activation (active > passive) in clusters comprising the left putamen, left insula, left thalamus, and lobule VIII of the right cerebellum. Reduced activation in the left insula and putamen was specific to own-hand feedback. Additionally, patients with SSD revealed reduced suppression (passive > active) in bilateral and medial parietal (including the right angular gyrus) and occipital areas, the right postcentral gyrus, cerebellum crus I, as well as the left medial superior frontal gyrus. Ego-disturbances were negatively correlated with left insula and putamen activation during active conditions, and with right angular gyrus activation patterns during passive conditions when own-hand feedback was presented.

These fMRI findings suggest that group differences are primarily evident during preparatory processes. Our results show that this preparatory neural activation is further linked to symptom severity, supporting the idea that the preparation of upcoming events as internal predictive mechanisms may underlie severe symptoms in patients with SSD. These findings could improve our understanding of other deficits in action planning, self-monitoring, and motor dysfunction in various psychiatric, neurological, and neurodegenerative disorders.

## Introduction

Impaired motor planning and movement disorder, disrupted executive functions, and psychotic symptoms, including hallucinations and delusions are prevalent across neurodegenerative, neurological, and psychiatric disorders. These disturbances are observed in patients with Alzheimer’s disease, Parkinson’s, cerebral palsy, dystonia, stroke, epilepsy, multiple sclerosis, amyotrophic lateral sclerosis, traumatic brain injury, and schizophrenia.^1–10^ Functional MRI (fMRI) studies suggested that movement and psychiatric disorders affect similar network connections between the basal ganglia and the cortex.^1^ This suggests a potential interface and shared interplay between movement and psychiatric disorders, despite differences in symptom severity, dynamics, and underlying mechanisms reported in neurological, neurodegenerative, and psychiatric disorders.^1,6,10,11^ Patients with schizophrenia can present with almost all these symptoms and impairments and are thus ideal candidates across these disorders to explore how movement preparation is reflected in the whole brain.

Core symptoms of schizophrenia spectrum disorders (SSD), such as auditory hallucinations and ego-disturbances (also referred to as “positive symptoms”) have been linked to impairments in action-feedback monitoring,^12–19^ reflected in aberrant efference copy model-based prediction of movement-related action consequences,^20–28^ sensory-motor feedback integration,^13,18,29–31^ perception of self-other distinction in the fine-tuned grip force and hand movement,^32–37^ sense of agency,^18,38–44^ and reduced neural suppression for self-generated (active) compared to externally generated (passive) hand movements.^18,45–50^ Although many studies have focused on action monitoring deficits in patients with SSD, neural differences between healthy subjects and patients with SSD regarding preparatory processes for hand movement effects remain except a few EEG studies^51–53^ mostly unexplored.

The execution of voluntary hand movements (active movement) requires internal preparation processes. These include the prediction of sensory action consequences, which must be maintained until movement initiation and ultimate execution. Regarding active movement intention and preparation, corollary discharge that enables own movement through motor command modulation, and effective internal sensory-motor prediction of movement feedback are often suggested to be generated as an efference copy of the motor command.^2,26,28,54,55^ This efference copy has been suggested to occur before the movement onset.^54,56–61^ In studies on sensory-motor specific predictive processes linked to efference-copy-based predictive processes, passive movements are frequently used as a control condition, since efference copy is not evident and thus assumed not to occur during passive movements.^22,50,55,59,62,63^

Movement preparation requires different cortical and sub-cortical area-specific balances of interactive neural pre-activation and inhibition of unwanted neural activation (pre-suppression).^57,64–66^ Patients with SSD, who may have cortical excitation and inhibition imbalance, especially in the motor-related areas, may deteriorate to show psychomotor abnormalities.^67,68^ Such a preparatory balanced neural function during movement preparation may further comprise adjustments in the internal prediction of subsequent movements and their consequences.^19,57,69–71^ Previous fMRI studies have suggested a potential engagement of the posterior medial frontal cortex, the pre-supplementary motor area, and the anterior cingulate cortex during the preparation of active eye and hand movements.^57^ A study that used continuous theta burst stimulation indicated that the supplementary motor area may relay the anticipated sensory consequences of a movement to the posterior sensory perceptual areas before reaching the primary motor cortex, thus before movement initiation.^65^ Recent delay adaptation studies highlighted how the hippocampus and superior frontal gyrus are linked to the recall of previous experiences regarding action outcome timing.^72,73^ Furthermore, a study that used magnetoencephalography to investigate reaching hand movements suggested that only the basal ganglia and cerebellum showed increased connectivity during movement. Interestingly, brain activity alterations and increased connectivity between most motor-related areas involved in motor control processes occurred before movement, whereas reports showed decreased neural activation in these regions during the movement execution.^58^ Although different in design, a study that compared self-paced to cued grasp movements further identified readiness blood-oxygen-level-dependent (BOLD) signals in brain areas, including the right inferior parietal lobe, right inferior frontal gyrus, bilateral supplementary motor area, bilateral precuneus, bilateral insula, bilateral visual cortex, and bilateral auditory cortex before movement onset.^66^ These findings potentially support our hypothesis that anticipatory neural coactivation processing in the engaged areas occurs during movement preparation, with perceptive processes taking place during the execution of a movement.

Regarding evidence of aberrant sensory-motor predictive processing, a study investigating the planning of gestures reported reduced neural activation in patients with SSD compared to healthy controls (HC) in the bilateral supplementary motor area, left superior temporal gyrus, left inferior occipital gyrus, left inferior parietal lobe, right middle frontal gyrus, and right inferior temporal gyrus, whereas stronger activations were recorded in the bilateral temporal pole.^14^ However, there are no fMRI studies manipulating action intention and preparation (i.e. active or passive) to investigate differences between patients with SSD and HC regarding sensory-motor preparatory processes before movement execution. Another notable research gap is how sensory-motor predictive processes prior to movement execution are affected by the identity of the feedback (e.g. Does the feedback display my own or someone else’s hand?). Lastly, researchers have yet to explore how differences in neural activation patterns existing prior to movement execution could be related to hallucinations and ego-disturbances in patients with SSD compared to HC. Previous EEG studies have revealed aberrant pre-movement motor preparation in SSD using the Bereitschaftspotential or Readiness Potential (RP) and Lateralized Readiness Potential (LRP), described as a potential marker of motor planning and initiation.^51–53^ However, it remains unclear how hand movement preparation processes are reflected in broader neural networks, which can be investigated using fMRI.

This study aimed to explore neural activation pattern differences between patients with SSD and HC regarding preparatory processes prior to movement execution by manipulating the initiation of the subsequent movement (i.e. active or passive movement) and visual feedback (i.e. feedback of one’s own or someone else’s hand). In a previous study using the same experimental setup and sample, we found neural activation differences between patients with SSD and HC during the execution of hand movements predominantly in the right angular gyrus, which have been linked to differences in self–other processing.^22^ Here, we extend these analyses to the pre-movement period to reveal group differences in pre-activation (i.e. active > passive) and pre-suppression (i.e. passive > active) effects.

## Materials and Methods

The participants, stimuli, and equipment of the current study have been detailed elsewhere.^22,62^ A summary of the essential aspects is given below.

### Participants

A total of 20 HC and 20 patients with SSD (19 with schizophrenia and one with schizoaffective disorder [SAD]) participating in the previously reported fMRI experiment^22^ were considered for the current analyses. Patients with SSD and HC were matched regarding their age, sex, and highest educational degree attained (see Table 1 for clinical and demographic characteristics). The patients with SSD were largely oligosymptomatic at the time of testing, as revealed by the Scale for the Assessment of Positive Symptoms (SAPS).^74^ The study was approved by the local ethics committee. Participants provided informed consent and were compensated for their participation.

**Table 1:** Demographic and clinical characteristics.

| Characteristics | HC (n = 20) |  | SSD (n = 20) |  |
| --- | --- | --- | --- | --- |
| Sex | 15 (male) | 5 (female) | 16 (male) | 4 (female) |
| Attained degree | 15 (LS), 5 (US), 2 (PS) |  | 13 (LS), 4 (US), 3 (PS) |  |
| Antipsychotic medication | None |  | 4 (none), 1 (FGA), 15 (SGA) |  |
| Age (Mean ± SD) | 38.4 ± 9.9 | | 38.5 ± 8.5 | |
| EHI score | 82.8 ± 29.3 | | 86.9 ± 17.5 | |
| MWT-B score | 27.1 ± 5.2 | | 27.8 ± 3.9 | |
| d2 score | 440.9 ± 85.6 | | 402.8 ± 103.1<sup>+</sup> | |
| TMT-A (seconds) | 31 ± 17 | | 29 ± 7 | |
| TMT-B (seconds) | 64 ± 29 | | 80 ± 31 | |
| WAIS FS score | 9.3 ± 1.7 | | 9.7 ± 1.4 | |
| WAIS BS score | 8.2 ± 1.7 | | 7.9 ± 1.6 | |
| SPQ-B: CP score | 0.9 ± 1.1 | | **3.8 ± 1.9** | |
| SAPS score | 1.9 ± 2.4 | | **12.1 ± 9.8** | |
| Hallucinations | 0.3 ± 0.9 | | 2 ± 4.2 | |
| Delusions | 0.4 ± 0.7 | | **5.6 ± 5.1** | |
| Delusions of reference | 0 ± 0 | | **1.1 ± 1.2** | |
| Delusions of being controlled | 0 ± 0 | | **0.5 ± 0.9** | |
| Residual positive symptoms | 1.2 ± 1.2 | | **4.2 ± 5.1** | |
| SANS score | 1.3 ± 2.0 | | **13.6 ± 12.2** | |

Note: LS: lower secondary; US: upper secondary; PS: post-secondary; EHI: Edinburgh Handedness Inventory; FGA: first-generation antipsychotics; SGA: second-generation antipsychotics; MWT-B: Mehrfachwahl-Wortschatz-Test (multiple choice vocabulary test); d2:d2 test of attention; TMT: Trail Making Test; WAIS: Wechsler Adult Intelligence Scale; FS: forward span; BS: backward span; SPQ-B: Schizotypal Personality Questionnaire-Brief; CP: Cognitive-Perceptual subscale; SAPS: Scale for the Assessment of Positive Symptoms; SANS: Scale for the Assessment of Negative Symptoms; HC: healthy control; SSD: schizophrenia spectrum disorder. Two patients were medicated with FGA as well as SGA; <sup>+</sup>missing data for one participant. Values and scores are the mean ± standard deviation. **Bold** values represent significant differences between HC and SSD patients ( $p < 0.05$ , uncorrected).

### Stimuli and Equipment

A custom-made MR-compatible passive movement device (PMD) was used. By gripping the handle of the PMD, the hand could be moved from the left (home position) to the right end and back to the left home position through a circular arc (central angle: ∼30 degrees; trajectory: about 5.5 cm, see Figure 1). Planning and hand movement can be self-induced or generated by the PMD using air pressure. Inside the MRI scanner, the PMD was placed beside the right thigh to facilitate right-hand movement. Handle position within ∼30° (movement trajectory) and direction of the hand movement were recorded via light emitting and detecting optical fibre cables integrated into the PMD.

**Figure 1.**
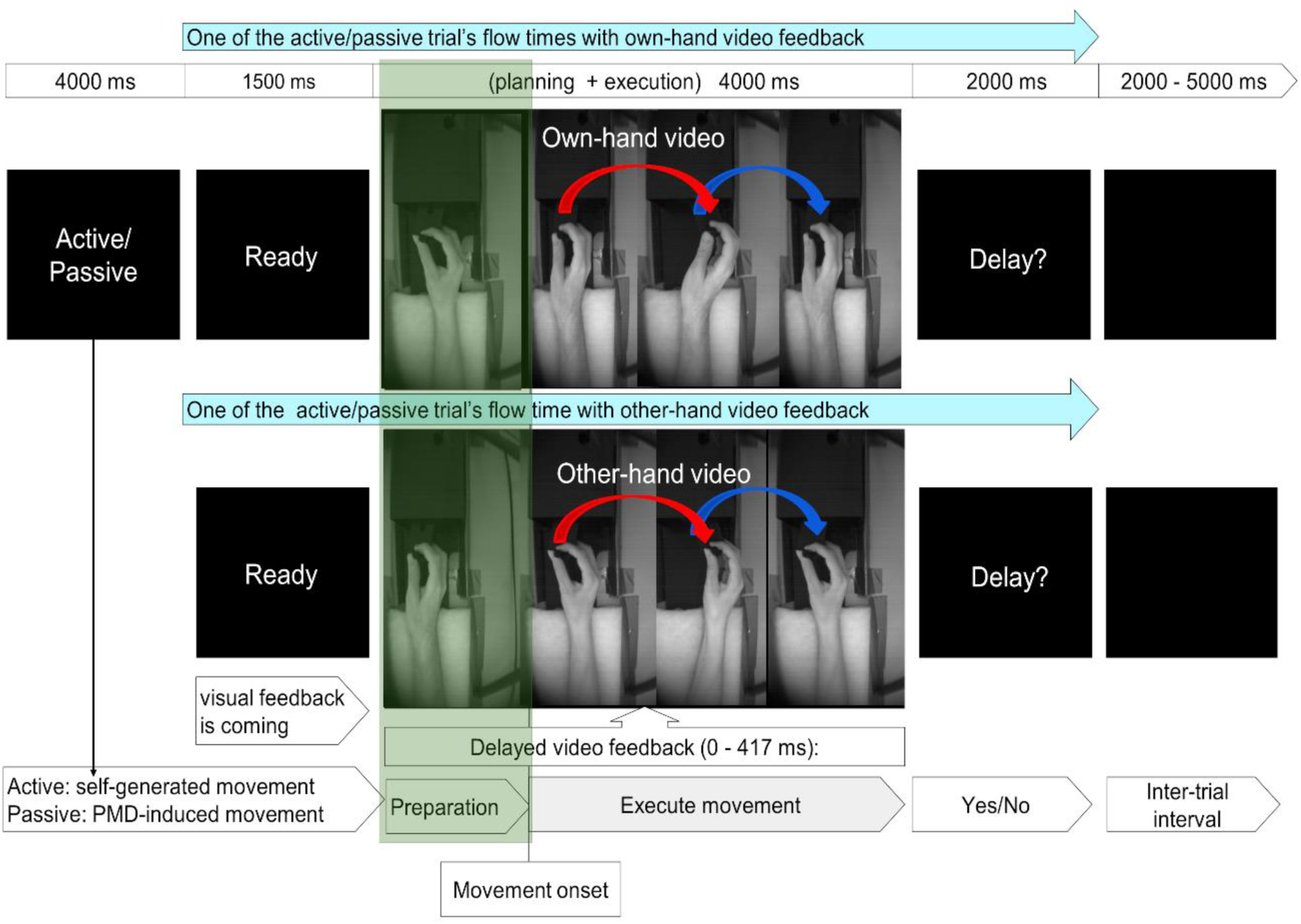
Experimental design. At the beginning of each run (48 trials), participants were instructed either to perform the hand movement during each trial themselves (“active” movements) or to relax their wrists and let their hands be moved by the PMD (“passive” movements). A trial commenced with a “Ready” cue, followed by visual feedback displaying the participant’s own or someone else’s hand. Subsequently, the participants were asked to report whether they noticed a delay (“Delay?”). A black screen with variable duration (2000– 5000 ms) was shown during the inter-trial interval. In the preparation period, the participant’s hand was at rest (1st frame marked with a green box), as the subject was instructed to move only to execute the hand movement. In the experiment, continuous video feedback was displayed, illustrated in four separate pictures to visualise the entire process of preparation and execution of the different movement directions. From the movement onset point, the hand moved from the left (2nd frame) to the right (3rd frame), and then the hand moved back to the left (shown by the 4th frame) position. Considering male participants for this figure, the upper row shows the sequence of a trial with “self” hand video feedback, while the lower row shows another trial with the “other” hand image from a female person. In the case of a female subject, a male hand image is displayed in the “other” hand video. The self-other hand was displayed randomly across the trials and runs. See the video demonstration available at http://doi.org/10.5281/zenodo.2621302.

During the experiment, video images of the participant’s hands were recorded via a high-speed camera (MRC High Speed, MRC Systems GmbH, Heidelberg, Germany; refresh rate: 4 ms) and displayed onto a screen (refresh rate: 60 Hz). Visual feedback from the monitor was shown on a tilted mirror inside the scanner. In 50% of the trials, a similarly positioned, pre-recorded hand of another person moving in accordance with the participant’s hand was displayed. During “other hand” trials, male participants were presented with the hand of a female person, and vice versa. Variable delays (0, 83, 167, 250, 334, and 417 ms + internal setup delay around 43 ms) were inserted between the actual hand movement and the video display. The participants task was to indicate whether the displayed hand movement in the video feedback was delayed by pressing a button using their left middle (yes) and index finger (no). These delays corresponded to the screen’s refresh rate (0, 5, 10, 15, 20, and 25 frames at 60 Hz). Custom-written software on a computer (Intel® Core™ i5-4,570 CPU, 3.20 GHz, 4 GB RAM, AMD Radeon HD8570D Graphics Card, 32-bit operating system, Windows 7 Professional [Microsoft Corporation, 2009]) was used to control the setup.

### Experimental Design

A mixed-factorial design with the within-subjects factors movement execution (active vs. passive) and video feedback (“self” vs. “other”), as well as a between-subjects factor group (patients with SSD vs. HC) was used, resulting in four conditions: self-active, self-passive, other-active, and other-passive.

Each run began with a cue (“Active” or “Passive”), instructing active or passive blocks of 24 trials each. Each trial was initiated with a “Ready” cue, followed by a randomised video display (duration: 4000 ms) of the hand (“self” or “other”) grasping the handle of the PMD. In active trials, the individual could freely perform the hand movement during the time the video was displayed, while in passive trials, the hand movement was programmed to be automatically initiated 500 ms (additionally, internal delay of compressor and PMD) after the camera/video onset by the PMD to maintain the similarity between the preparation and movement onset timing in active and passive trials. Both in active and passive condition, equally and randomly distributed 6 different delays within 0 - 417 ms were applied between the video feedback and the actual hand movement. Subsequently, a “Delay?” cue appeared, and the participants had to report if they noticed a delay or not. Trials concluded with a black screen (inter-trial interval, randomised duration between 2000-5000 ms).

### Procedure

There was a behavioural preparatory session to familiarise the participants with the setup. During the preparatory session, the participants sat upright in front of a computer screen. The preparatory session was followed by an fMRI session with two runs of 48 trials each. Patients who were either unable or unwilling to be in the MRI scanner (n = 18) performed two behavioural sessions outside the scanner (see Uhlmann et al.^22^) and were not considered in the analysis.

In the fMRI experiment, the subjects were in a supine position inside the MRI, while the PMD was placed next to the right thigh. The participants were instructed to maintain their grip of the PMD handle with their right hand, using the index finger and thumb for the upper part and the remaining fingers for the lower part of the handle. Hand movements consisted of an extension from the left to the right and a contraction to move it back from the right to the left home position. The participants were asked to complete a movement in about 1500 ms to make it consistent with the passive condition, which was trained in the preparatory session. At the end of the experiment, all subjects filled out a post-experiment questionnaire.

### Functional Data Acquisition

Data were acquired using a 3 T Magnetom Trio Tim scanner (Siemens, Erlangen, Germany) with a 12-channel head coil. For functional data, a T2*-weighted gradient-echo echoplanar imaging sequence (TR: 1650 ms, TE: 25 ms, flip angle: 70°) was procured. In each of the two runs, 330 volumes were collected in descending order, each covering 34 axial brain slices (matrix: 64 × 64, field of view [FoV]: 192 mm × 192 mm, slice thickness: 4 mm, voxel size: 3 mm × 3 mm × 4.6 mm [with a 0.6 mm gap]). Anatomical images were acquired using a T1-weighted MPRAGE sequence (TR: 1900 ms, TE: 2.26 ms, flip angle: 9°) with a matrix of 256 × 256, FoV of 256 mm × 256 mm, slice thickness of 1 mm, and voxel size of 1 mm × 1 mm × 1.5 mm [including a 0.5 mm gap].

### Imaging Data Preprocessing

The standard procedures from Statistical Parametric Mapping (SPM12 version 7, Wellcome Trust Centre for Neuroimaging, University College London, UK) were implemented in MATLAB 2017a (Mathworks Inc.) to analyse the fMRI data. The preprocessing steps involved realignment, coregistration between anatomical and functional images, segmentation, normalisation to the Montreal Neurological Institute (MNI) standard space (resampled to a voxel size of 2 mm × 2 mm × 2 mm), and smoothing (8 mm × 8 mm × 8 mm full-width at half-maximum Gaussian kernel). The framewise displacement (FD) between consecutive volumes was computed. Only < 10% of all FD values exceeded 1 mm in any run. Here, the focus was on the fMRI contrast relevant to the preparation before movement onset versus the movement execution period (Figure 1).

### Statistical Analyses

All analyses related to BOLD signal were done using statistical parametric mapping (SPM) integrated into MATLAB 2017a. Correlation related analyses were performed using JASP (Jeffreys’s Amazing Statistics Program; version 0.18.3; JASP team, 2024). In the analyses, the acquired BOLD response between the camera onset and camera offset from each trial was included, while a hand was displayed on the screen during each experimental condition. This BOLD signal was segmented into preparation-based BOLD and movement execution-based BOLD responses. For the preparation, this included the time from the camera onset to the movement onset (average duration for self-active: HC = 654 ms, standard deviation [SD] = 219, SSD = 747 ms, SD = 355; for self-passive: HC = 908 ms, SD = 90, SSD = 887 ms, SD = 85; other-active: HC = 663 ms, SD = 228, SSD = 727 ms, SD = 288; and other-passive: HC = 907 ms, SD = 83, SSD = 885 ms, SD = 82). For movement execution, we considered the time from movement onset to the camera offset (average duration for self-active: HC = 3356 ms, SD = 219, SSD = 3261 ms, SD = 355; for self-passive: HC = 3102 ms, SD = 90, SSD = 3121 ms, SD = 84; other-active: HC = 3346 ms, SD = 229, SSD = 3281 ms, SD = 288; and other-passive: HC = 3103 ms, SD = 83, SSD = 3123 ms, SD = 81). In general, SSD patients showed no significant differences in preparation duration compared to HC. In addition, no group interaction effect was found in repeated measure ANOVA. Therefore, for each participant, 4 preparation and 4 movement execution regressors were modelled as conditions of interest, and cue and question periods as well as the 6-motion parameter were included as nuisance regressors in the general linear model. The regressors were convolved with the canonical haemodynamic response function. To remove low-frequency noise from the time series, a 128 s high-pass filter was applied. Lastly, we contrasted regressors of interest against the implicit baseline for each participant, and the resulting contrast estimates were entered into a group-level full factorial model.

To correct for multiple comparisons at *p* < 0.05, we applied the identical cluster threshold of 104 voxels and cluster defining threshold of *p* < 0.005 uncorrected, as previously determined by Monte Carlo simulations.^22^ (Note: family-wise error [FWE] cluster-corrected *p* values are also provided in Tables 2 and 3) This ensures consistency with the previous analyses and results. Coordinates are listed using the standard MNI152 space. To explore activation in preparation periods, we analysed the following T-contrasts within HC and SSD groups and compared them in conjunction analyses as well as in between-group contrast: prep(active>passive); contrast for the commonalities: HC prep(active>passive) Ո SSD prep(active>passive); contrast for the differences: HC prep(active>passive) > SSD prep(active>passive). The same analyses were conducted for passive>active contrast. For the condition and group-specific effects, contrast masking was implemented. To investigate the group-specific effects, we analysed HC prep(active>passive) > SSD prep(active>passive) masked by HC prep(active>passive); SSD prep(active>passive) > HC prep(active>passive) masked by SSD prep(active>passive); HC prep(passive>active) > SSD prep(passive>active) masked by HC prep(passive>active). For condition-specific effects, we analysed HCprep((selfact-selfpas)-(otheract-otherpas))>SSDprep((selfact-selfpas)-(otheract-otherpas)) masked by HC prep (self-active).

**Table 2:**
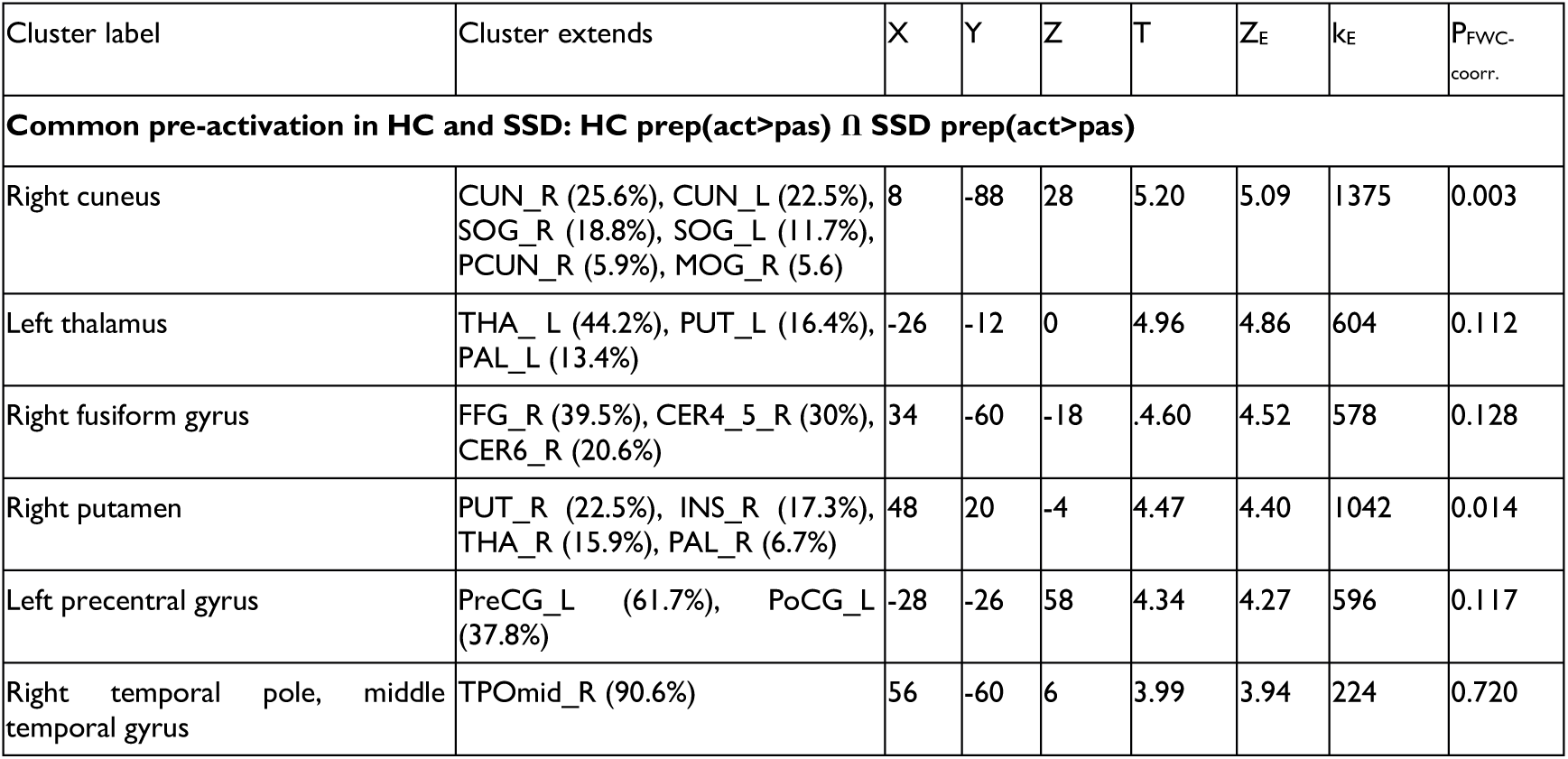
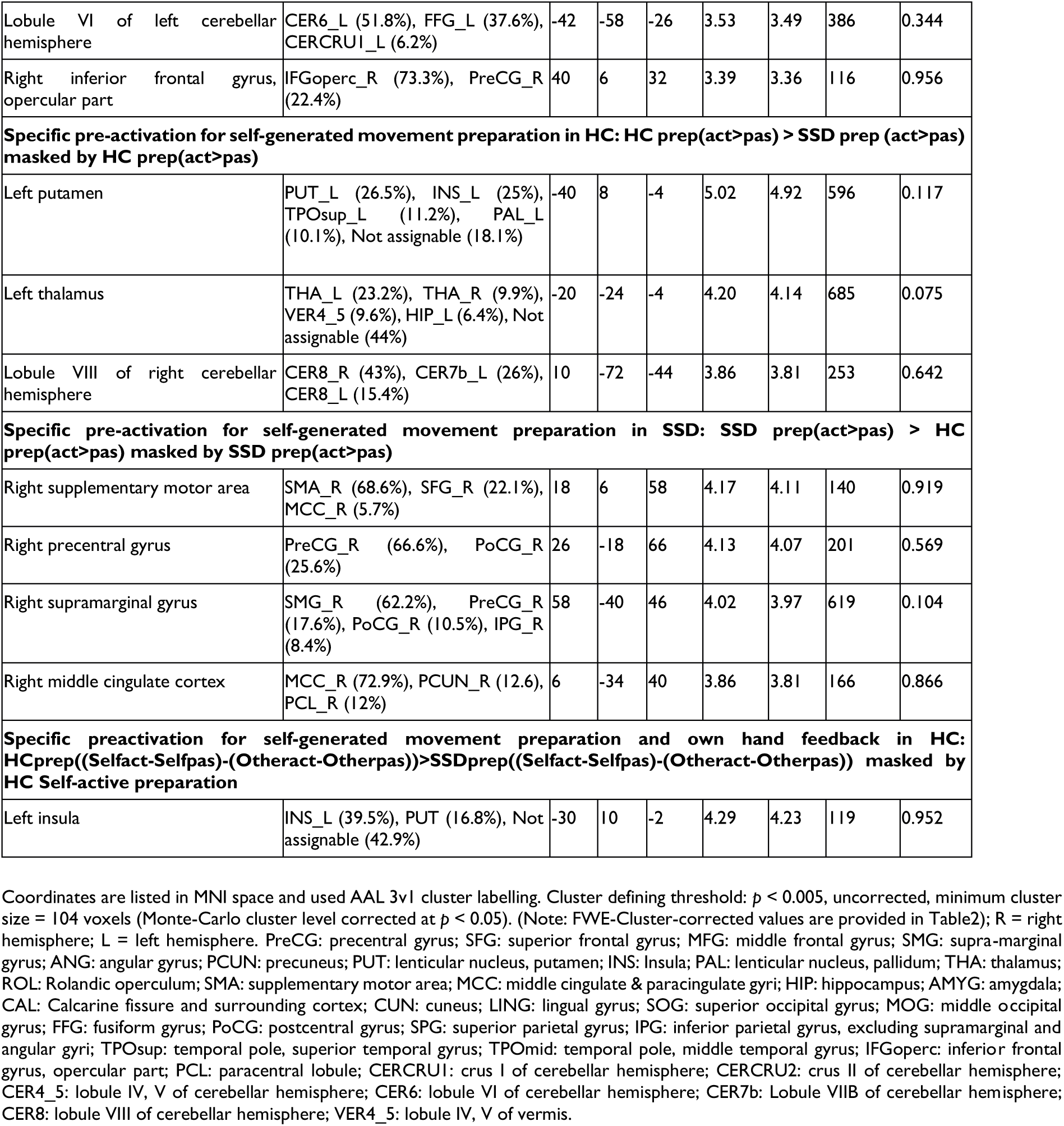
Results regarding pre-activation (active > passive)

| Cluster label | Cluster extends | X | Y | Z | T | Z<sub>E</sub> | k<sub>E</sub> | P<sub>FWC-coorr.</sub> |
| --- | --- | --- | --- | --- | --- | --- | --- | --- |
| **Common pre-activation in HC and SSD: HC prep(act>pas) ∩ SSD prep(act>pas)** |  |  |  |  |  |  |  |  |
| Right cuneus | CUN_R (25.6%), CUN_L (22.5%), SOG_R (18.8%), SOG_L (11.7%), PCUN_R (5.9%), MOG_R (5.6) | 8 | -88 | 28 | 5.20 | 5.09 | 1375 | 0.003 |
| Left thalamus | THA_L (44.2%), PUT_L (16.4%), PAL_L (13.4%) | -26 | -12 | 0 | 4.96 | 4.86 | 604 | 0.112 |
| Right fusiform gyrus | FFG_R (39.5%), CER4_5_R (30%), CER6_R (20.6%) | 34 | -60 | -18 | 4.60 | 4.52 | 578 | 0.128 |
| Right putamen | PUT_R (22.5%), INS_R (17.3%), THA_R (15.9%), PAL_R (6.7%) | 48 | 20 | -4 | 4.47 | 4.40 | 1042 | 0.014 |
| Left precentral gyrus | PreCG_L (61.7%), PoCG_L (37.8%) | -28 | -26 | 58 | 4.34 | 4.27 | 596 | 0.117 |
| Right temporal pole, middle temporal gyrus | TPOmid_R (90.6%) | 56 | -60 | 6 | 3.99 | 3.94 | 224 | 0.720 |
| Lobule VI of left cerebellar hemisphere | CER6_L (51.8%), FFG_L (37.6%), CERCRUI_L (6.2%) | -42 | -58 | -26 | 3.53 | 3.49 | 386 | 0.344 |
| Right inferior frontal gyrus, opercular part | IFGoperc_R (73.3%), PreCG_R (22.4%) | 40 | 6 | 32 | 3.39 | 3.36 | 116 | 0.956 |
| **Specific pre-activation for self-generated movement preparation in HC: HC prep(act>pas) > SSD prep (act>pas) masked by HC prep(act>pas)** |  |  |  |  |  |  |  |  |
| Left putamen | PUT_L (26.5%), INS_L (25%), TPOsup_L (11.2%), PAL_L (10.1%), Not assignable (18.1%) | -40 | 8 | -4 | 5.02 | 4.92 | 596 | 0.117 |
| Left thalamus | THA_L (23.2%), THA_R (9.9%), VER4_5 (9.6%), HIP_L (6.4%), Not assignable (44%) | -20 | -24 | -4 | 4.20 | 4.14 | 685 | 0.075 |
| Lobule VIII of right cerebellar hemisphere | CER8_R (43%), CER7b_L (26%), CER8_L (15.4%) | 10 | -72 | -44 | 3.86 | 3.81 | 253 | 0.642 |
| **Specific pre-activation for self-generated movement preparation in SSD: SSD prep(act>pas) > HC prep(act>pas) masked by SSD prep(act>pas)** |  |  |  |  |  |  |  |  |
| Right supplementary motor area | SMA_R (68.6%), SFG_R (22.1%), MCC_R (5.7%) | 18 | 6 | 58 | 4.17 | 4.11 | 140 | 0.919 |
| Right precentral gyrus | PreCG_R (66.6%), PoCG_R (25.6%) | 26 | -18 | 66 | 4.13 | 4.07 | 201 | 0.569 |
| Right supramarginal gyrus | SMG_R (62.2%), PreCG_R (17.6%), PoCG_R (10.5%), IPG_R (8.4%) | 58 | -40 | 46 | 4.02 | 3.97 | 619 | 0.104 |
| Right middle cingulate cortex | MCC_R (72.9%), PCUN_R (12.6%), PCL_R (12%) | 6 | -34 | 40 | 3.86 | 3.81 | 166 | 0.866 |
| **Specific preactivation for self-generated movement preparation and own hand feedback in HC: HCprep((Selfact-Selfpas)-(Otheract-Otherpas))>SSDprep((Selfact-Selfpas)-(Otheract-Otherpas)) masked by HC Self-active preparation** |  |  |  |  |  |  |  |  |
| Left insula | INS_L (39.5%), PUT (16.8%), Not assignable (42.9%) | -30 | 10 | -2 | 4.29 | 4.23 | 119 | 0.952 |

Coordinates are listed in MNI space and used AAL 3v1 cluster labelling. Cluster defining threshold: $p < 0.005$ , uncorrected, minimum cluster size = 104 voxels (Monte-Carlo cluster level corrected at $p < 0.05$ ). (Note: FWE-Cluster-corrected values are provided in Table2); R = right hemisphere; L = left hemisphere. PreCG: precentral gyrus; SFG: superior frontal gyrus; MFG: middle frontal gyrus; SMG: supra-marginal gyrus; ANG: angular gyrus; PCUN: precuneus; PUT: lenticular nucleus, putamen; INS: Insula; PAL: lenticular nucleus, pallidum; THA: thalamus; ROL: Rolandic operculum; SMA: supplementary motor area; MCC: middle cingulate & paracingulate gyri; HIP: hippocampus; AMYG: amygdala; CAL: Calcarine fissure and surrounding cortex; CUN: cuneus; LING: lingual gyrus; SOG: superior occipital gyrus; MOG: middle occipital gyrus; FFG: fusiform gyrus; PoCG: postcentral gyrus; SPG: superior parietal gyrus; IPG: inferior parietal gyrus, excluding supramarginal and angular gyri; TPOsup: temporal pole, superior temporal gyrus; TPOmid: temporal pole, middle temporal gyrus; IFGoperc: inferior frontal gyrus, opercular part; PCL: paracentral lobule; CERCRUI: crus I of cerebellar hemisphere; CERCRU2: crus II of cerebellar hemisphere; CER4\_5: lobule IV, V of cerebellar hemisphere; CER6: lobule VI of cerebellar hemisphere; CER7b: Lobule VIIb of cerebellar hemisphere; CER8: lobule VIII of cerebellar hemisphere; VER4\_5: lobule IV, V of vermis.

**Table 3:** Results regarding pre-suppression (passive > active)

| Cluster label | Cluster extend | X | Y | Z | T | Z<sub>E</sub> | k<sub>E</sub> | P<sub>FWC-coorr.</sub> |
| --- | --- | --- | --- | --- | --- | --- | --- | --- |
| **Group differences in pre-suppression: HC prep (pas > act) > SSD prep (pas > act)** |  |  |  |  |  |  |  |  |
| Right postcentral gyrus | PoCG_R (12.9%), IPG_R (11.5%), SMG_R (10.3%), PreCG_R (9.2%), SFG_R (6.5%), ANG_R (6.3%), PCUN_L (5.8%), Not assignable (15.5%) | 46 | -54 | -42 | 4.59 | 4.51 | 4826 | < 0.001 |
| Left supramarginal gyrus | SMG_L (64%), IPG_L (25.3%), PoCG_L (7.5%) | -64 | -34 | 28 | 3.92 | 3.87 | 292 | 0.542 |
| Left supplementary motor area | SMA_L (71.6%), PCL_L (20.8%), SFG_L (6.6%) | -8 | -12 | 70 | 3.81 | 3.76 | 197 | 0.791 |
| Right lingual gyrus | LING_R (40.4%), LING_L (32.6%), CAL_R (18.9%), CAL_L (6.6%) | 10 | -68 | 0 | 3.79 | 3.74 | 381 | 0.352 |
| Crus I of right cerebellar hemisphere | CERCRUI_R (28%), CERCRU2_L (22.4%), CERCRUI_L (16.2%), CER6_R (13.5%), CERCRU2_R (9.9%), VER7 (9.4%) | 12 | -76 | -24 | 3.68 | 3.63 | 192 | 0.884 |
| Right calcarine | CAL_R (60.3%), Not assignable (33.9%) | 28 | -72 | 4 | 3.45 | 3.41 | 174 | 0.848 |
| Left superior parietal gyrus | SPG_L (84.4%), IPG_L (15.7%) | -30 | -60 | 54 | 3.42 | 3.39 | 115 | 0.957 |
| SFGmedial_L | SFGmedial_L (43.1%), SFG_L (35.8%), SMA_L (9.5%) | -12 | 18 | 50 | 3.34 | 3.38 | 137 | 0.924 |
| **HC prep(pas>act) ∩ SSD prep(pas>act): common suppression in HC and SSD** |  |  |  |  |  |  |  |  |
| Crus II of right cerebellar hemisphere | CERCRU2_R (76.7%), CERCRUI_R (23.3%) | 24 | -76 | -36 | 4.02 | 3.96 | 197 | 0.791 |
| **Group differences in areas specifically where HC show pre-suppression: HC prep(pas>act) > SSD prep (pas>act) masked by HC prep(pas>act)** |  |  |  |  |  |  |  |  |
| Right inferior parietal gyrus | IPG_R (28%), ANG_R (16.6%), SPG_R (9.9%), PCUN_R (7.3%), SMG_R (5.7%), Not assignable (31.4%) | 46 | -54 | 42 | 4.59 | 4.51 | 807 | 0.041 |
| Left precuneus | PCUN_L (37.5%), SPG_L (7%), CUN_L (5%), Not assignable (43.9) | -14 | -60 | 38 | 4.08 | 4.02 | 510 | 0.182 |
| Left supramarginal gyrus | SMG_L (95.8%) | -64 | -34 | 28 | 3.92 | 3.87 | 154 | 0.952 |
| Right lingual gyrus | LING_R (83.8%), CAL_R (14.3%) | 10 | -68 | 0 | 3.79 | 3.74 | 154 | 0.539 |
| Right postcentral gyrus | PoCG_R (79.1%), PCUN_R (10.6%), SPG_R (9.8%) | 26 | -42 | 70 | 3.78 | 3.73 | 254 | 0.639 |
| Crus I of right cerebellar hemisphere | CERCRUI_R (30.6%), CERCRU2_L (21.2%), CER6_R (14%), CERCRUI_L (14%), CERCRU2_R (11.2%) | 12 | -76 | -24 | 3.68 | 3.63 | 170 | 0.857 |
| Right calcarine | CAL_R (58.7%), Not assignable (35.3%) | 28 | -72 | 4 | 3.45 | 3.41 | 167 | 0.864 |
| Left medial superior frontal gyrus | SFGmedial_L (43.2%), SFG_L (36.4%), SMA_L (8.3%), Not assignable (12.1%) | -12 | 18 | 50 | 3.45 | 3.41 | 167 | 0.864 |
| **Specific pre-suppression in SSD: SSD prep(pas>act) > HC prep (pas>act) masked by SSD prep(pas>act)** |  |  |  |  |  |  |  |  |
| Left middle temporal gyrus | MTG_L (32.5%), ITG_L (18.5%), Not assignable (44.4%) | -52 | -10 | -24 | 3.42 | 3.38 | 135 | 0.927 |
| **Group differences between preparation and movement-based activation:**<br>**[HC prep (act > pas) > mov (act > pas)] > [SSD prep (act > pas) > mov (act > pas)]** |  |  |  |  |  |  |  |  |
| Left putamen | PUT_L (33.2 %), INS_L (22 %), STG_L (15.5 %), PAL_L (10.1 %), not assignable (14.1 %) | -40 | 8 | -4 | 3.34 | 3.21 | 277 | 0.579 |

Coordinates are listed in MNI space and used AAL 3v1 cluster labelling. Cluster defining threshold: $p < 0.005$ , uncorrected, minimum cluster size = 104 voxels (Monte-Carlo cluster level corrected at $p < 0.05$ ) (Note: FWE-cluster-corrected values are provided additionally in Table 3); R = right hemisphere; L = left hemisphere. PreCG: precentral gyrus; SFG: superior frontal gyrus; SFGmedial: superior frontal gyrus, medial; MFG: middle frontal gyrus; SMG: supra marginal gyrus; ANG: angular gyrus; PCUN: precuneus; PUT: lenticular nucleus, putamen; INS: insula; PAL: lenticular nucleus, pallidum; THA: thalamus; ROL: Rolandic operculum; SMA: supplementary motor area; MCC: middle cingulate & paracingulate gyri; PCC: posterior cingulate gyrus; AMYG: amygdala; CAL: calcarine fissure and surrounding cortex; CUN: cuneus; LING: lingual gyrus; FFG: fusiform gyrus; PoCG: postcentral gyrus; SPG: superior parietal gyrus; IPG: inferior parietal gyrus, excluding supramarginal and angular gyri; TPOsup: temporal pole: superior temporal gyrus; TPOmid: temporal pole, middle temporal gyrus; IFGoperc: inferior frontal gyrus, opercular part; PCL: paracentral lobule; STG: superior temporal gyrus; MTG: middle temporal gyrus; ITG: inferior temporal gyrus; CERCRUI: crus I of cerebellar hemisphere; CERCRU2: crus II of cerebellar hemisphere; CER4\_5: Lobule IV, V of cerebellar hemisphere; CER6: lobule VI of cerebellar hemisphere; VER7: Lobule VII of vermis.

The cluster label assumes the name of the brain area with the highest percent contribution to the cluster (using AAL 3v1 cluster labelling).^75^ The cluster extension brain regions (>5% contribution) and their contribution percentages are listed in Tables 2 and 3. Only clusters from which more than 50% of voxels could be labelled are listed. A cluster label with an MNI152 space coordinate (X Y Z) was used to interpret the results. Thus, all of the results are shown here in the MNI152 space’s coordinate.

### Exploratory Correlation Analyses

In particular, negative symptoms, ego-disturbances, and hallucinations have been related to dysfunctional self-other discrimination and the prediction of one’s own movement consequences.^51,76–80^ Therefore, we investigated the correlation of positive symptoms, assessed using related subscales of hallucinations (items 1–7), delusions of reference (item 14), delusions of being controlled (item 15)), and residual positive symptoms (items 21–35) of the assessment of positive symptoms (SAPS),^74^ with neural activations (extracted eigenvariates) of areas showing specific pre-activation in active (left insula/putamen, lobule VIII of right cerebellar hemisphere) or passive (right angular gyrus, left middle temporal gyrus) conditions with own hand video feedback. The same areas neural activations correlations with negative symptoms were assessed using related subscales of affective flattening or blunting (items 1–8), alogia (items 9–13), avolition/apathy (items 14–17), anhedonia/asociality (items 18–22), and attention (items 23–25 of the scale for the assessment of negative symptoms (SANS).^81^ To explore specificity regarding the positive symptoms, partial correlations were performed by partialling out total SANS score (see Supplementary Table 1). Analyses were performed using JASP (Jeffreys’s Amazing Statistics Program; version 0.18.3; JASP team, 2024).

### Data availability

The data information supporting this study’s findings is published in the following repository link. https://doi.org/10.5281/zenodo.14858692

## Results

### fMRI Results

#### Activation during the pre-movement period (pre-activation effects; active > passive)

We found activation for active compared to passive conditions during the pre-movement period both in HC (Figure 2A) and patients with SSD (Figure 2B). Conjunction analyses confirmed an overlap of pre-activation effects in subcortical (predominantly bilateral thalamus, putamen), cortical (predominantly cuneus and right frontal operculum), and cerebellar brain regions (Figure 2C, Table 2). Group differences were evident as a reduced pre-activation in the left putamen, left thalamus, and lobule VIII of the right cerebellar hemisphere area (Figure 2D, Table 2) and increased pre-activation in the right supplementary motor area, right precentral gyrus, right supramarginal gyrus, right middle cingulate gyrus (Figure 2E, Table 2), and right angular gyrus (Figure 3C) in patients with SSD compared to HC.

**Figure 2.**
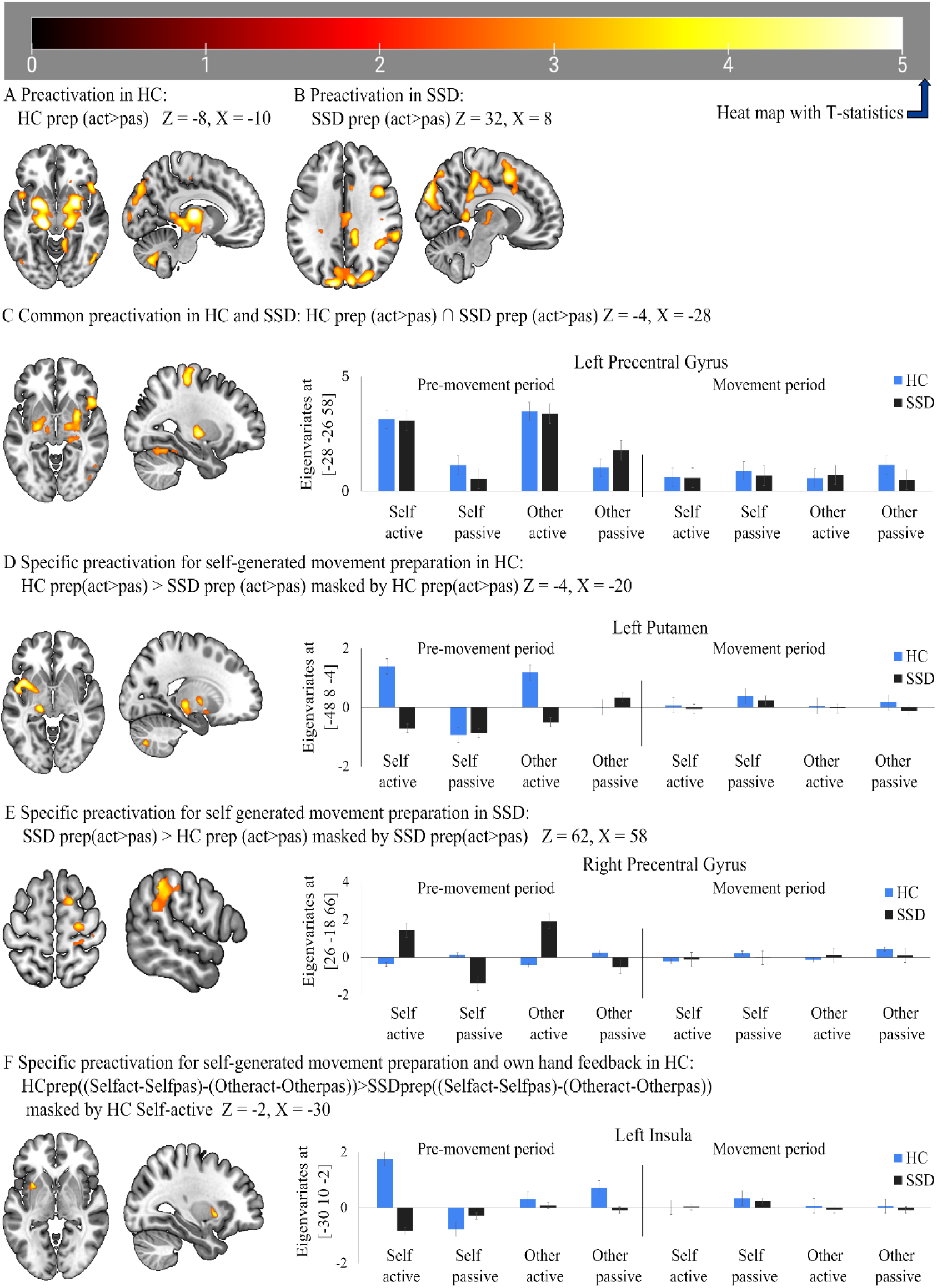
Neural activation in active compared to passive condition. (**A**) activation in healthy control (HC) at Z = -8, X = -10; (**B**) activation in schizophrenia spectrum disorder (SSD) at Z = 32, X = 8; (**C**) common activated brain areas between HC and SSD at Z = -4, X = -28; (**D**) activation specific for HC at Z = -4, X = -20; (**E**) activation specific for SSD at Z = 62, X = 58; (**F**) differentiable activated brain area between HC and SSD at Z = -2, X = -30. The performed statistical test is a T-contrast with T-values ranging from 1 to 5. HC: healthy control, (n= 20); SSD: schizophrenia spectrum disorder, (n = 20).

**Figure 3.**
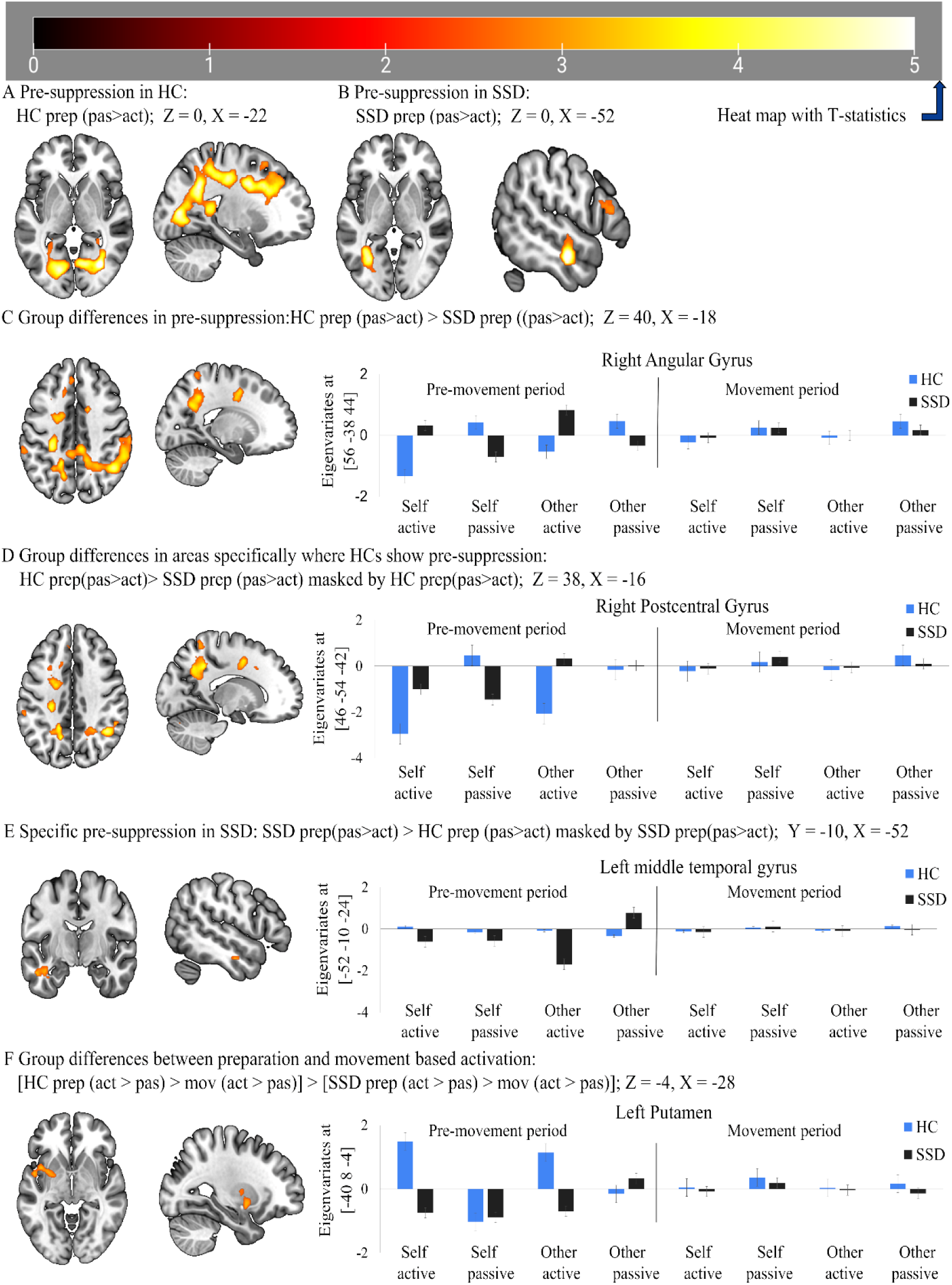
Neural activation in passive compared to active conditions. (**A**) activation in healthy control (HC) at Z = 0, X = -22; (**B**) activation in schizophrenia spectrum disorder (SSD) at Z = 0, X = -52; (**C**) differentiable activated brain areas between HC and SSD at Z = 32, X = -18; (**D**) activation specific for HC at Z = 38, X = -16; (**E**) pre-suppression specific for SSD at Z = -24, X = -52; **(F)** group-specific differences of activation pattern during preparation than movement execution shown at Z = -4, X = -28. The performed statistical test is a T-contrast with T-values ranging from 1 to 5. HC: healthy control, (n= 20); SSD: schizophrenia spectrum disorder, (n = 20).

Lastly, we found a specific pre-activation in the left insula for the own-hand video feedback condition in healthy subjects, which was significantly reduced in patients with SSD (see Figure 2F, Table 2), as revealed by an interaction analysis of the factors self/other, active/passive, and group.

#### Pre-suppression effect during the pre-movement period (pre-suppression effects; passive > active)

We found activation for passive compared to active conditions during the pre-movement period in both HC (Figure 3A) and patients with SSD (Figure 3B). Conjunction analyses revealed a significant overlap of pre-suppression [neural inhibition or reduced neural activation for active compared to passive movements (passive>active) reflected by negative eigenvariates during movement preparation] effects at crus II of the right cerebellar hemisphere (Table 3). Group differences were evident in a reduced pre-suppression effect in the right inferior parietal gyrus, left precuneus, left supramarginal gyrus, right lingual gyrus, right postcentral gyrus, crus I of the right cerebellum, right calcarine fissure and surrounding cortex, and left medial superior frontal gyrus regions (Figure 3D, Table 3) in patients with SSD compared to HC. Lastly, we found a specific pre-suppression in the left middle temporal gyrus, which was significantly increased in patients with SSD (see Figure 3E, Table 3).

### Movement period

Concerning commonalities between preparation and movement, we found no significant cluster, either for activation or for suppression. Intriguingly, group differences between preparation and movement-based activation (a three-factor interaction of group [HC > SSD], condition [active > passive], and phase [preparation > movement]) patterns revealed a cluster comprising mainly left putamen and left insula (Table 3, Figure 3F). These findings indicate that compared to patients with SSD, HC exhibited higher brain activation primarily during the preparation phase, whereas no significant differences were observed during movement execution. This could further reflect that preparation-induced brain activation patterns are different from movement execution-induced areas.

### Exploratory correlations with symptoms

The preparatory neural activation in the left insula/putamen for active hand movement with feedback from the own hand revealed a negative correlation to delusions of reference, delusions of being controlled, residual positive symptoms (Figure 4A), avolition/apathy, and anhedonia/asociality. The preparatory neural activation in the right angular gyrus for passive hand movement with own-hand video feedback was negatively correlated with hallucinations (Figure 4B) and affective flattening or blunting. Analyses with negative symptoms thus suggest that these correlations were not specific to positive symptoms i.e. hallucinations and ego-disturbances, but also correlated to negative symptoms i.e. affective flattening or blunting, avolition/apathy, and anhedonia/asociality. Additionally, partial correlation by partialling out the total score of SANS (scale for the assessment of negative symptoms) showed a similar result (see Supplementary Table 1 and Table 2), indicating quite independent impairments related to positive and negative symptoms.

**Figure 4.**
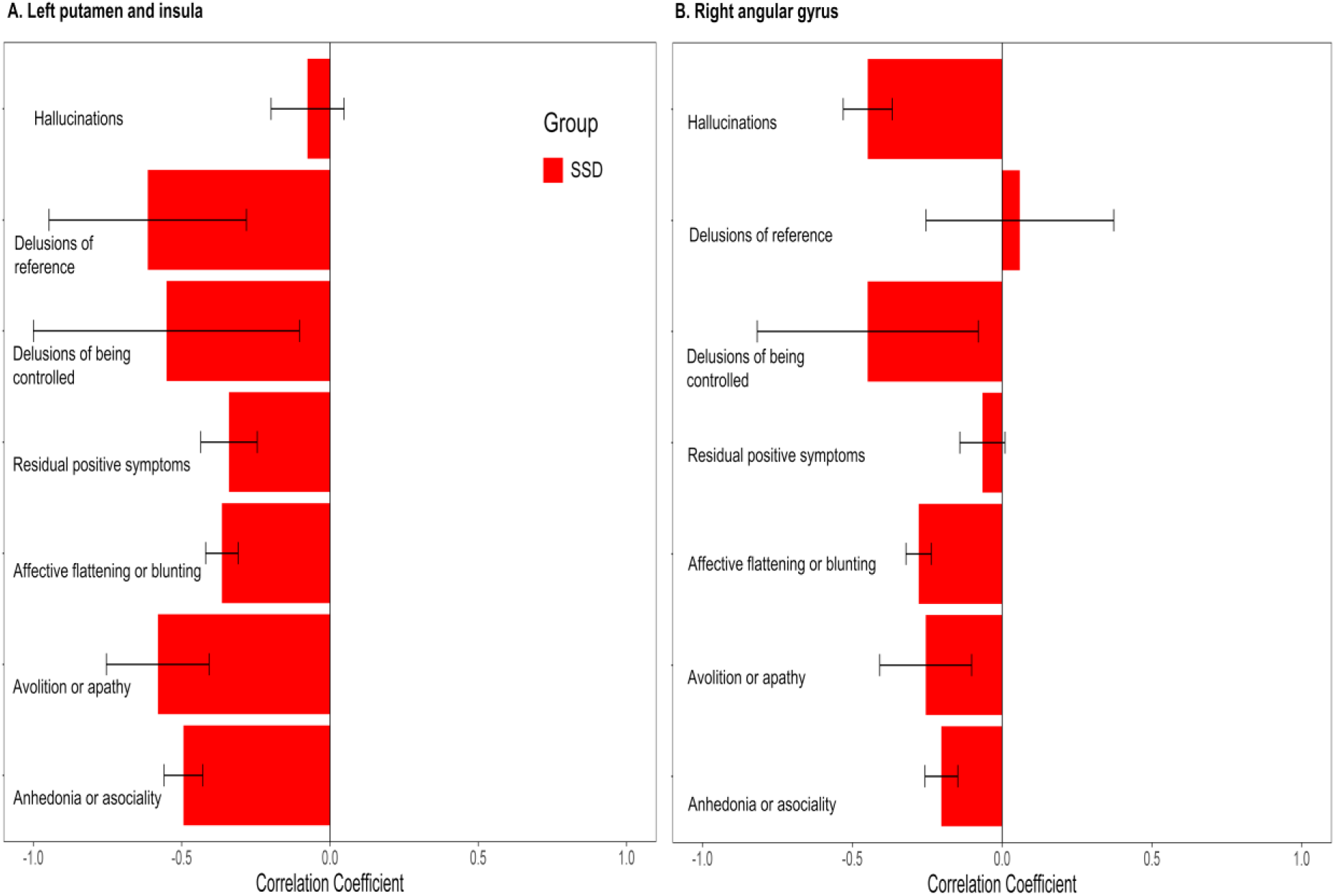
Exploratory correlation of neural activity with positive and negative symptoms. (**A**) Pre-activation in the left putamen and insula for active conditions with own hand feedback (see Figure 2F). (**B**) Pre-activation in the right angular gyrus for passive conditions with own hand feedback. Hallucinations (SAPS questions 1–7), delusions of reference (SAPS question 14), delusions of being controlled (SAPS question 15), and residual positive symptoms (SAPS questions 21–35), affective flattening or blunting (SANS items 1–8), avolition/apathy (SANS items 14–17), and anhedonia/asociality (SANS items 18–22). The error bar represents the standard error mean and reflects a 95% confidence interval calculated from the Pearson correlation coefficient.

## Discussion

To our knowledge, this is the first fMRI study revealing group differences in the pre-movement period, specifically focusing on planning and preparation for the processing of one’s own hand feedback in active compared to passive hand movement conditions. Here, we demonstrated for the first time with fMRI that patients with SSD differ from healthy controls as early as the pre-movement period, extending previous EEG findings on premovement related potentials (e.g., Bereitschaftspotential) in schizophrenia.^51–53^ Our SSD cohort exhibited less activation in the left insula and putamen and less suppression in the parietal brain regions (including the right angular gyrus) before movement. We further showed that the feedback-specific reduced neural activation of the left insula and putamen before movement execution in patients with SSD is associated with ego-disturbances (delusions of reference and delusions of being controlled), indicating impairments in the preparation period as a potential basis for the reduced sense of agency or passivity symptoms in these patients.

Previous fMRI research has mainly focused on action–outcome monitoring deficits in patients with SSD. While EEG studies in SSD, although limited to electrode over motor areas and not the hand movement, findings of reduced LRP amplitude and delayed LRP onset before index finger press/thumb movement described as disrupted response selection, planning, and preparation; while the absence of such difference during response execution implicates deficit during response preparation in SSD.^82^ Similarly, reduced or abolished preparatory RP amplitude described in schizophrenia (n=9, no female) with left and right thumb-button press, ^83^ during motor preparation of figure copying task,^84^ during single and sequential finger-button press tasks manipulating speed in SSD (n = 11, four male) highlighted a global preparatory planning dysfunction reflected by reduced amplitude and delayed RP.^53^ Thus, support that motor abnormalities in SSD are inherent ^29,85–90^ and more consistently found during the planning and preparation phase rather than in execution. Although the relevance of predictive mechanisms for action feedback processing is widely acknowledged, preparatory effects before movement have rarely been explored in previous well-controlled fMRI studies. In this study, the pre-activation of the insula was specific to the video feedback of one’s own hand in healthy subjects and significantly reduced in SSD patients. These aberrant pre-activation patterns may contribute to the previously reported deficits in action-outcome monitoring.^22^

We found neural activation during active hand movement preparation revealed in conjunction analysis comparing the distributed regions of HC and patients with SSD, including the left precentral gyrus, the right middle temporal gyrus, lobule VI of the left cerebellar hemisphere, left cuneus, and right putamen. Thus, for the first time, we showed that pre-activation in these areas seems to be intact or preserved in patients with SSD. Higher neural activation in the preparation for active compared to passive movement in these areas may emphasise that action preparation involves many brain areas beyond the sensory-motor brain regions. It further demonstrates that movement feedback processing is not restricted to a sensory-motor attenuation perspective but can go along with pre-movement activation enhancement.

However, we also found group differences in pre-activation and pre-suppression of neural activity in the pre-movement period. Higher neural activation among healthy subjects (Table 2, Figure 2D) revealed clusters primarily consisting of the left putamen and insula, left thalamus, and lobule VIII of the right cerebellar hemisphere area during active preparation compared to patients with SSD. This pattern may reflect altered functional preparation for one’s own actions and their consequences, potentially leading to a deficit in self-other differentiation in SSD patients. This interpretation is in line with the finding that the left putamen regulates right-hand movement preparation and execution, and it co-activates during motor tasks.^91^ Furthermore, the putamen, along with other striatal areas, receives input from the cortex, filters, synchronises, and integrates them to drive a focused output to the left insula and left thalamus for further control of cortical and cerebellar activity.^91,91–95^

We found specific pre-activation of active movement with own-hand feedback preparation. This is evident in the left insula and putamen in active preparation (Table 2, Figure 2D), and in a subcluster specifically for the active hand movement preparation with own hand feedback (Table 2, Figure 2F). This suggests that the activation of the putamen and insula is specifically relevant for the preparation of self-generated hand-movement feedback. Aberrant left insula activation may thus reflect impairment of these functions and facilitate the deficit of multisensory function in the self-consciousness and action awareness in general,^96,97^ body ownership, and the sense of agency.^98–101^ The insula has also been implicated in conscious retrospective error awareness,^19^ abnormal prior-based calibration,^102,103^ recalibration of prediction effect,^19,57^ and deficit in hosting interoceptive prediction by comparing outcomes and updating preparation upon error finding.^104^ Therefore, it is likely that the pre-activation of insula and putamen for self-generated hand movement feedback reflects an action outcome predictive process specific to one’s own hand feedback. Further, we observed that the left insula and putamen showed a negative correlation of pre-activation with delusions of reference, delusions of being controlled, avolition/apathy, and anhedonia/asociality (Figure 4A) (Supplementary Table 1), further demonstrating that less pre-activation goes along with more ego-disturbances and more avolition/apathy as well as anhedonia/asociality. Thus, reduced neural activation of the left insula and putamen during feedback-specific preparation in patients with SSD is associated with both positive symptoms (e.g., delusions of reference and delusions of being controlled) and negative symptoms, indicating impairments in the preparation period as a potential basis for reduced sense of agency or passivity symptoms in these patients. While exploratory partial correlation analyses indicate some symptom specific effects (Supplementary Table 2), strong conclusions about symptom specific effects cannot be drawn from this exploratory study due to the small sample size.

Regarding suppression in the premovement period, our result shows that BOLD suppression (passive > active) for self-generated movement feedback was already present during movement preparation. Notably, this pre-suppression was reduced in patients with SSD in the right angular gyrus (Figure 3C) and medial superior frontal gyrus (Table 3), with suppression instead of activation in the passive condition, indicating impaired function of the right angular gyrus and left medial superior frontal gyrus during preparation. The insula is suggested as a supramodal area, processing multi-sensory signals from the body.^105,106^ It integrates sensory signals in the context of emotion and motivation^98^ and provides impetus to the dorsomedial frontal cortex to initiate and sustain movement, potentially bridging body awareness and movement.^25,107^ This active movement preparation initiation in the left medial superior frontal gyrus potentially drives suppression on the right angular gyrus, which is activated on the preparation for passive movement, potentially driving attention and readiness towards the externally generated movement. These roles of the medial superior frontal gyrus and right angular gyrus are supported by a meta-analysis,^108^ connectivity and transcranial magnetic stimulation studies on self-agency,^109–111^ and studies using transcranial direct current stimulation. These findings suggest that following the dorsolateral prefrontal cortex (this could be the left medial superior frontal gyrus in our findings), the right angular gyrus could act as a circuit breaker to detect and decouple self-other actions.^112,113^ Thus, our finding extends previously reported agency impairments related to the right angular gyrus in SSD^22^ and highlights its primary relevance in the preparation period as a starting point of BOLD suppression. Regarding the right angular gyrus, we found a negative correlation of pre-activation in passive conditions with hallucinations and affective flattening or blunting (Figure 4B) (Supplementary Table 1). Thus, patients with SSD demonstrating less pre-activation in passive conditions showed more hallucinations and affective flattening or blunting. These findings also align with prior EEG studies demonstrating associations between impaired movement preparation and negative symptoms in schizophrenia.^51,114–116^ These studies highlight how reduced RP (Bereitschaftspotential) reflects deficits in motor planning, which may contribute to avolition and psychomotor slowing. Similarly, our results suggest that reduced neural activation during movement preparation could underlie broader impairments in motor function and negative symptomatology in SSD.

In group differences (Table 2, Figure 2D), HC showed significantly higher pre-activation in lobule VIII of the right cerebellar hemisphere area than in patients with SSD. This may indicate that lobule VIII of the right cerebellar hemisphere area is impaired in SSD. This raises the question of how the impaired function of lobule VIII of the right cerebellar hemisphere could reveal the underlying deficit in preparatory processing and the prediction of the consequences of hand movements. Along this line, fMRI studies and reviews on overt finger tapping movement often suggest that lobules VI and VIII of the right cerebellar hemisphere are the hosts of the sensorimotor cerebellar area.^114–117^ Inadequate activation of lobule VIII of the right cerebellar hemisphere may reflect the primary impaired cerebellar area underlying hand movement preparation and self-other agency distinction. We found a negative correlation in the lobule VIII of the right cerebellar hemisphere, similar to the insula and putamen, and linked increased delusions of reference and avolition/apathy to less pre-activation. Furthermore, the thalamus is a central hub that engages in the filtering, integrating, and calibrating of the received signal before relaying through cortico-thalamus-cerebellum-cortical networks during movement planning and execution; however, these coordinated networks are often suggested to be disrupted in SSD.^20,118–121^ Therefore, our finding of aberrant thalamic activation (Table 2, Figure 2D) may further indicate reduced coordination in SSD patients during preparation.

Regarding the correlation of symptoms and motor preparation, patients with SSD revealed distinct patterns of EEG abnormalities from HC. A study (n = 31, 11 female, two-left handed) using thumb opposition and flexion tasks found reduced early RP and late RP amplitudes and movement-related motor potential peak at the central (Cz) electrode, were negatively correlated to the total SANS score.^51^ Similarly, reduced late LRP amplitude at C4 is linked to negative symptoms, while reduced early RP amplitude is linked to patients with positive and mixed symptoms.^122^ Another study of brisk right fist closure in patients with early course schizophrenia (n=10, one female) found a correlation between reduced RP at C1 and weaker post-movement event-related synchronisation (ERS) with the high examination of anomalous self-experience (EASE) score; negative correlation of reduced RP and higher positive symptoms assessed by positive and negative syndrome scale (PANS) score; indicating disrupted interaction between impaired preparatory RP, post-movement ERS, and self-disorder including self-awareness and delusions of being controlled.^80^ In our findings, a similar effect of positive and negative symptoms on preparatory neural activation was observed (Supplementary Table 1). Avolition and apathy are recognised as core negative symptoms^79^ which might underly the reduction in the initiation and persistence of goal-directed activities, disrupting movement preparatory neural activation pattern. Together our findings complement prior EEG studies on pre-movement preparation in SSD by providing evidence of altered activation patterns in subcortical and cerebellar regions during this phase. While EEG studies have primarily focused on electrophysiological markers such as the Bereitschaftspotential, our fMRI results highlight the involvement of specific brain regions, including the left insula and putamen, which may contribute to impairments in self-monitoring and agency.

Finally, in SSD patients, we observed specific hyper neural activation in the right precentral gyrus, right supramarginal gyrus, and right middle cingulate cortex, especially in active conditions (see Table 2, Figure 2E), which may indicate an abnormal additional brain activation pattern, potentially reflecting aberrant or mixed hemispheric lateralisation. These hyperactivations in the right hemisphere instead of the left hemisphere may reflect the tendency to engage alternative or complementary brain areas and strategies in active hand movement preparation. Furthermore, our results of suppression in the left middle temporal gyrus among patients with SSD (Table 3, Figure 3E) may support the reduced volume and thickness of the left middle temporal gyrus underlying reduced or distributed cerebral asymmetry of the hemisphere in schizophrenia patients and their relatives, as suggested by many studies.^76,123–125^

Taken together, reduced activation of the left insula, putamen, and lobule VIII of the right cerebellar hemisphere during active conditions, along with a lack of suppression in an active and over-suppression instead of activation during the passive condition in the right angular gyrus, may precursory of inadequate preparation for action outcome monitoring during movement execution in SSD patients. Furthermore, the negative correlation observed between symptoms and activation patterns suggests that this mechanism may be involved in ego-disturbances and hallucinations.

## Limitations

The relatively small sample size needs to be considered a potential limitation regarding the generalizability of the study’s results, especially the correlation analyses. The application of statistical thresholds like family-wise error correction and correction for multiple comparisons has led to the disappearance of significance in most areas. In future experiments, large samples with separate preparation and execution trials should be investigated. Furthermore, exploratory analyses suggest that correlations are not limited to hallucinations and ego-disturbances, as negative symptoms or residual positive symptoms are also related to pre-activation patterns. Therefore, we cannot exclude the fact that the general symptom load is related to altered neural processes in the pre-movement period. Future studies with larger cohorts are needed to validate these associations and further disentangle the relationships between neural activation patterns, positive symptoms, and negative symptoms

## Conclusion

This study investigated activation and suppression patterns before right-hand movement execution and for the first time, showed commonalities and differences in these preparatory processes between HC and patients with SSD. Based on our fMRI results, the deficits in action– outcome monitoring in schizophrenia manifest in the preparation period on a neural level before an action is executed. The negative correlation of both positive and negative symptoms with preparatory neural activation patterns further indicates that impaired preparatory neural processes are likely linked to clinically relevant features, such as ego-disturbances, hallucinations, avolition/apathy, anhedonia/asociality, and affective flattening or blunting. Thus, impaired preparatory processes could lead to difficulties in attributing feedback to the movement of one’s own hand, potentially underly ego-disturbances, hallucinations, avolition/apathy, anhedonia/asociality, and affective flattening or blunting in schizophrenia. However, given the exploratory nature of our correlation analyses and the small sample size, these results should be interpreted cautiously. Considering these fMRI findings, we propose that disturbances exist during preparation for the movement, rather than in its execution, resulting in impairment in patients with SSD. Our findings may reflect a broader, similarly disrupted cortico-cerebellar functional area that could exist across neurological and neurodegenerative disorders. Future research could use fMRI and diffusion tensor imaging to map commonalities, uniquely impaired areas, and disrupted structural and functional connectivity in movement preparation across schizophrenia, neurodegenerative, and neurological disorders. This approach would identify shared and disorder-specific underlying modulatory cortical areas that could be targeted with specific interventions to improve motor dysfunctions, executive functions, hallucinations, and ego-disturbances.

## Acknowledgements

We thank the Core Facility Brain Imaging Marburg and Lukas Uhlmann, Mareike Pazan, Anastasia Benedyk, Volker Besmens, Laila Noor, Lars Schwenzer, Jens Sommer, Olaf Steinsträter, and Dominik Vaughan for data collection as well as Jens Sommer for assistance with the implementation of the experimental setup.

## Supplementary material

See attached files

