## Supplementary Table for "Aberrant preparation of hand movement in schizophrenia spectrum disorder: An fMRI study"

### Supplementary material to the manuscript:
### Aberrant preparation of hand movement in schizophrenia spectrum disorder: An fMRI study

**Table 1: Correlation between neural activation (cluster's eigenvariates) and symptom score in SSD patients**

| Cluster consisting mainly | symptom | Pearson correlation coeeficient: r | p | Effect size (Fisher's z) | Spearman correlation coeeficient: rho | p | Effect size (Fisher's z) |
|---|---|---|---|---|---|---|---|
| Left insula and putamen<br><br>preparation for active movement with own hand feedback | SAPS_I | -0.076 | 0.752 | -0.076 | -0.281 | 0.230 | -0.289 |
| | SAPS_14 | -0.614 | **0.004** | -0.716 | -0.585 | 0.007 | -0.669 |
| | SAPS_15 | -0.473 | **0.035** | -0.514 | -0.466 | 0.039 | -0.504 |
| | residual positive symptoms | -0.341 | 0.142 | -0.355 | -0.278 | 0.241 | -0.282 |
| | SANS_I | -0.076 | 0.752 | -0.076 | -0.281 | 0.230 | -0.289 |
| | SANS_II | -0.241 | 0.305 | -0.246 | -0.282 | 0.228 | -0.290 |
| | SANS_III | -0.580 | **0.007** | -0.663 | -0.608 | 0.004 | -0.706 |
| | SANS_IV | -0.494 | **0.027** | -0.542 | -0.510 | 0.022 | -0.563 |
| | SANS_V | 0.069 | 0.774 | 0.069 | 0.0.099 | 0.877 | 0.100 |
| Left middle temporal gyrus<br><br>preparation for passive movement with own hand feedback | SAPS_I | -0.495 | **0.027** | -0.542 | -0.31 | 0.183 | -0.321 |
| | SAPS_14 | -0.087 | 0.716 | -0.087 | -0.155 | 0.515 | -0.156 |
| | SAPS_15 | -0.468 | **0.037** | -0.508 | -0.307 | 0.187 | -0.318 |
| | residual positive symptoms | -0.162 | 0.496 | -0.163 | -0.090 | 0.707 | -0.090 |
| | SANS_I | -0.495 | **0.027** | -0.542 | -0.310 | 0.183 | -0.321 |
| | SANS_II | 0.046 | 0.847 | 0.046 | -0.117 | 0.624 | -0.117 |
| | SANS_III | -0.270 | 0.250 | -0.277 | -0.151 | 0.526 | -0.152 |
| | SANS_IV | 0.029 | 0.905 | 0.029 | -0.0008 | 0.997 | -0.0008 |
| | SANS_V | 0.235 | 0.319 | 0.239 | 0.259 | 0.271 | 0.265 |
| Right angular gyrus<br><br>preparation for passive movement with own hand feedback | SAPS_I | -0.448 | **0.047** | -0.483 | -0.502 | 0.024 | -0.551 |
| | SAPS_14 | 0.059 | 0.804 | 0.059 | 0.079 | 0.742 | 0.079 |
| | SAPS_15 | -0.402 | 0.079 | -0.426 | -0.085 | 0.720 | -0.086 |
| | residual positive symptoms | -0.066 | 0.783 | -0.066 | -0.029 | 0.903 | -0.029 |

| | | | | | | | |
|---|---|---|---|---|---|---|---|
| | SANS_I | -0.448 | **0.047** | -0.483 | -0.502 | 0.024 | -0.551 |
| | SANS_II | -0.118 | 0.620 | -0.119 | -0.222 | 0.348 | -0.225 |
| | SANS_III | -0.255 | 0.278 | -0.261 | -0.089 | 0.710 | -0.089 |
| | SANS_IV | -0.203 | 0.391 | -0.206 | -0.241 | 0.307 | -0.246 |
| | SANS_V | 0.257 | 0.274 | 0.263 | 0.298 | 0.201 | 0.308 |
| Lobule VIII of right cerebellar hemisphere<br><br>preparation for active movement with own hand feedback | SAPS_I | 0.038 | 0.874 | 0.038 | -0.177 | 0.456- | -0.179 |
| | SAPS_14 | -0.616 | **0.004** | -0.719 | -0.609 | 0.004 | -0.707 |
| | SAPS_15 | -0.300 | 0.198 | -0.310 | -0.492 | -0.028 | -0.538 |
| | residual positive symptoms | -0.447 | **0.048** | -0.480 | -0.116 | 0.627 | -0.116 |
| | SANS_I | 0.038 | 0.874 | 0.038 | -0.177 | 0.458 | -0.179 |
| | SANS_II | -0.287 | 0.220 | -0.295 | -0.417 | 0.067 | -0.444 |
| | SANS_III | -0.453 | **0.045** | -0.488 | -0.491 | 0.028 | -0.537 |
| | SANS_IV | -0.374 | 0.105 | -0.393 | -0.485 | 0.030 | -0.529 |
| | SANS_V | -0.599 | **0.005** | -0.691 | -0.378 | 0.100 | -0.398 |

Note: SAPS scale for the assessment of positive symptoms, SAPS_I hallucinations, SAPS_II delusions, SAPS_14 delusions of reference, SAPS_15 delusions of being controlled, SAPS_III bizarre behavior, SAPS_IV positive formal thought disorder, SAPS_res residual positive symptom (SAPS_III + SAPS_IV + SAPS_V). SPQ-B: schizotypal personality questionnaire-brief, SPQ-B_cogn_perc SPQ-B cognitive perceptual deficit, SPQ_Interpers SPQ interpersonal deficit, SPQ_desorg SPQ-B disorganization, SPQ_total SPQ-B total score accumulated from SPQ_cogn_perc + SPQ_interpers + SPQ_desorg, SANS scale for the assessment of negative symptoms, SANS_I affective flattening or blunting, SANS_II alogia, SANS_III avolition/apathy, SANS_IV anhedonia/asociality, SANS_V attention, SANS_summe_global sum of item 22 to 24, SANS_Gesamtkalenwert sum of SANS_I to SANS_V. **Bold** values represent significant differences between HC and SSD patients ($p < 0.05$, uncorrected).

**Table 2: Partial correlation (partialling out SANS score) between neural activation (cluster's eigenvariates) and symptom score in SSD patients**

| Cluster consisting mainly | symptom | Pearson correlation coeeficient: r | p | Effect size (Fisher's z) | Spearman correlation coeeficient: rho | p | Effect size (Fisher's z) |
|---|---|---|---|---|---|---|---|
| Left insula and putamen<br><br>preparation for active movement with own hand feedback | SAPS_1 | 0.258 | 0.286 | 0.264 | 0.011 | 0.964 | 0.011 |
| | SAPS_14 | -0.471 | **0.042** | -0.512 | -0.414 | 0.078 | -0.440 |
| | SAPS_15 | -0.321 | 0.195 | -0332 | -0.348 | 0.158 | -0.364 |
| | residual positive symptoms | -0.245 | 0.312 | -0.250 | -0.222 | 0.361 | -0.226 |
| Left middle temporal gyrus<br><br>preparation for passive movement with own hand feedback | SAPS_1 | -0.475 | **0.040** | -0.517 | -0.291 | 0.226 | -0.300 |
| | SAPS_14 | 0.046 | 0.850 | 0.046 | -0.094 | 0.702 | -0.094 |
| | SAPS_15 | -0.516 | **0.028** | -0.571 | -0.436 | 0.071 | -0.467 |
| | residual positive symptoms | -0.112 | 0.648 | -0.113 | -0.069 | 0.790 | -0.069 |
| Right angular gyrus<br><br>preparation for passive movement with own hand feedback | SAPS_1 | -0.332 | 0.165 | -0.345 | -0.374 | 0.115 | -0.393 |
| | SAPS_14 | 0.364 | 0.125 | 0.382 | 0.430 | 0.066 | 0.460 |
| | SAPS_15 | -0.295 | 0.235 | -0.304 | 0.251 | 0.314 | 0.257 |
| | Sa_SAPS_15 | -0.520 | **0.027** | -0.570 | -0.368 | 0.133 | -0.386 |
| | residual positive symptoms | 0.034 | 0.890 | 0.034 | 0.039 | 0.873 | 0.039 |
| Lobule VIII of right cerebellar hemisphere<br><br>preparation for active movement with own hand feedback | SAPS_1 | 0.384 | 0.107 | 0.405 | 0.045 | 0.855 | 0.045 |
| | SAPS_14 | -0.489 | **0.033** | -0.535 | -0.520 | 0.020 | -0.588 |
| | SAPS_15 | 0.052 | 0.839 | 0.052 | -0.435 | 0.071 | -0.466 |
| | residual positive symptoms | -0.232 | 0.340 | -0.236 | 0.113 | 0.646 | 0.113 |

Note: SAPS scale for the assessment of positive symptoms, SAPS_I hallucinations, SAPS_II delusions, SAPS_14 delusions of reference, SAPS_15 delusions of being controlled, SAPS_III bizarre behavior, SAPS_IV positive formal thought disorder, SAPS_res residual positive symptom (SAPS_III + SAPS_IV + SAPS_V). SPQ-B: schizotypal personality questionnaire-brief, SPQ-B_cogn_perc SPQ-B cognitive perceptual deficit, SPQ_Interpers SPQ interpersonal deficit, SPQ_desorg SPQ-B disorganization, SPQ_total SPQ-B total score accumulated from SPQ_cogn_perc + SPQ_interpers + SPQ_desorg, SANS scale for the assessment of negative symptoms, SANS_I affective flattening or blunting, SANS_II alogia, SANS_III avolition/apathy, SANS_IV anhedonia/asociality, SANS_V attention, SANS_summe_global sum of item 22 to 24, SANS_Gesamtkalenwert sum of SANS_I to SANS_V. Bold values represent significant differences between HC and SSD patients ($p < 0.05$, uncorrected
